# Longitudinal Assessment of SARS-CoV-2 Specific T Cell Cytokine-Producing Responses for 1 Year Reveals Persistence of Multi-Cytokine Proliferative Responses, with Greater Immunity Associated with Disease Severity

**DOI:** 10.1101/2022.01.18.476864

**Authors:** Jonah Lin, Ryan Law, Chapin S. Korosec, Christine Zhou, Wan Hon Koh, Mohammad Sajjad Ghaemi, Philip Samaan, Hsu Kiang Ooi, FengYun Yue, Anne-Claude Gingras, Antonio Estacio, Megan Buchholz, Patti Lou Cheatley, Katerina Pavinski, Samira Mubareka, Allison J. McGeer, Jerome A. Leis, Jane M. Heffernan, Mario Ostrowski

## Abstract

Cellular-mediated immunity is critical for long-term protection against most viral infections, including coronaviruses. We studied 23 SARS-CoV-2-infected survivors over a one year post symptom onset (PSO) interval by *ex vivo* cytokine ELISpot assay. All subjects demonstrated SARS-CoV-2-specific IFN-γ, IL-2, and Granzyme B (GzmB) T cell responses at presentation, with greater frequencies in severe disease. Cytokines, mainly produced by CD4+ T cells, targeted all structural proteins (Nucleocapsid, Membrane, Spike) except Envelope, with GzmB > IL-2 > IFN-γ. Mathematical modeling predicted that: 1) cytokine responses peaked at 6 days for IFN-γ, 36 days for IL-2, and 7 days for GzmB, 2) severe illness was associated with reduced IFN-γ and GzmB, but increased IL-2 production rates, 3) males displayed greater production of IFN-γ, whereas females produced more GzmB. *Ex vivo* responses declined over time with persistence of IL-2 in 86% and of IFN-γ and GzmB in 70% of subjects at a median of 336 days PSO. The average half-life of SARS-CoV-2-specific cytokine-producing cells was modelled to be 139 days (∼4.6 months). Potent T cell proliferative responses persisted throughout observation, were CD4 dominant, and were capable of producing all 3 cytokines. Several immunodominant CD4 and CD8 epitopes identified in this study were shared by seasonal coronaviruses or SARS-CoV-1 in the Nucleocapsid and Membrane regions. Both SARS-CoV-2-specific CD4+ and CD8+ T cell clones were able to kill target cells, though CD8 tended to be more potent.

**Importance:** Our findings highlight the relative importance of SARS-CoV-2-specific GzmB-producing T cell responses in SARS-CoV-2 control, shared CD4 and CD8 immunodominant epitopes in seasonal coronaviruses or SARS-CoV-1, and indicate robust persistence of T cell memory at least one year after infection. Our findings should inform future strategies to induce T cell vaccines against SARS-CoV-2 and other coronaviruses.

## Introduction

As of November 2021, the World Health Organization (WHO) had reported over 250 million confirmed cases and more than 5 million deaths due to COVID-19 (1). SARS-CoV-2 Spike antibody inducing vaccines have dramatically slowed the infection and death rate in the developed world, however, breakthrough infections due to delta and other variant viruses continue, and infection rates continue to rise in non-vaccinated regions (2-7).

Neutralizing antibodies against SARS-CoV-2 Spike are a major effector in protection against infection and disease. However, some level of protection has been observed in vaccinated individuals prior to serum neutralizing antibody development, indicating other arms of the immune system likely play a role (8). Also, individuals with recent other beta coronavirus infections who do not have cross-reactive antibodies appear to have more limited disease after SARS-CoV-2 (9), suggesting other mediators of protection other than antibodies. T cells are generally important for viral clearance and disease protection, whereby CD4+ T cells can enhance antibody maturation and CD8+ T cell-mediated killing of infected cells. In regard to SARS-CoV-2 infection, a potent early T cell response correlates with disease outcome (10) and T cell responses are induced during convalescence (11-13), which have been shown to persist for up to 8 months (14).

A number of important questions remain regarding the role of T cells in COVID-19 immunity and disease. The role of CD4+ versus CD8+ T cell memory in disease and protection are unclear. The relative importance of various T cell effector cytokines such as IFN-γ, IL-2, and granzyme B are still poorly understood. It is suspected that mapping of CD4+ and CD8+ T cell epitopes and with attention to regions outside of Spike protein may inform the next generation of SARS-CoV-2 vaccines, given the emergence of antibody escape variants in vaccinated persons. It is still unclear how long T cell memory can persist in SARS-CoV-2 infection, given that re-infection is very common with coronaviruses in general (15, 16), albeit often with reduced severity. Such information should in the future reveal how T cells operate as correlates of protection against infection and disease.

In the current study, we prospectively followed a cohort of SARS-CoV-2 infected individuals with varying levels of disease outcome by using highly sensitive cytokine ELISpot assays to follow SARS-CoV-2 antigen-specific cytokine responses averaging about one year of follow up. We modelled the kinetics of the T cell cytokine response with time. In addition, we mapped a panel of epitopes to structural proteins that characterized dominant T cell responses in this cohort and show effector function and killing capabilities of both CD4+ and CD8+ T cell clones.

## Methods

### Human Subjects and Study Approval

Healthy subjects and individuals with COVID-19 infection as diagnosed by positive nasopharyngeal SARS-CoV-2 PCR were recruited and informed consent was obtained from for blood draws and/or leukapheresis through an REB approved protocol (St. Michael’s Hospital/Unity Health Toronto REB20-044). Asymptomatic COVID-19 negative controls had no history of viral infection and negative serology (IgG) for SARS-CoV-2 full Spike, Spike receptor binding domain, and Nucleocapsid proteins by ELISA as previously described by us (17). All human subjects research was done in compliance with the Declaration of Helsinki.

### PBMC Isolation

Whole blood or leukapheresis samples were acquired from St. Michael’s Hospital, Unity Health (Toronto, CA). Peripheral blood mononuclear cells (PBMC) were isolated by centrifugation using Ficoll-Paque PLUS (GE Healthcare). PBMCs were resuspended in R10 media consisting of RPMI-1640 (Wisent), 10% heat-inactivated fetal bovine serum (FBS; Wisent), 10 mM HEPES (Wisent), 2 mM L-glutamine (Wisent), and 100 U penicillin-streptomycin solution (Wisent). PBMCs were diluted 1:1 with freezing medium consisting of 20% DMSO (Sigma-Aldrich) in FBS and aliquoted for −150°C storage. PBMCs used in the T cell epitope mapping were sent to Scisco Genetics Inc. (Seattle, WA) for HLA typing **(Table S1).**

### SARS-CoV-2 Peptides and Peptide Pools Synthesis

SARS-CoV-2 15-mer peptides overlapping by 11 amino acids from the four main structural proteins of the SARS-CoV-2 reference sequence (NC_045512.2) were synthesized by Genescript (Piscataway, New Jersey, USA) for T cell epitope mapping. In total, 212 individual peptides containing 102 Nucleocapsid (N), 12 Envelope (E), 49 Membrane (M), and 49 Spike (S) 15-mers were synthesized. The S 15-mers used for the T cell epitope mapping spanned only the receptor-binding domain (RBD) and transmembrane domain (TM) due to cost considerations. Using these peptides, 30 matrix peptide pools containing 19-23 15-mers were generated using the “Deconvolute This!” program Version 1.0 (18, 19). For longitudinal assessment of participants’ cytokine responses, 4 peptide master pools for each structural protein (N, E, M, S-RBD + S-TM) and 2 peptide master pools containing the full S protein (S1 and S2) were synthesized by Genescript and JPT (Berlin, Germany), respectively.

### *Ex Vivo* ELISpot Assay

*Ex vivo* ELISpot assays were performed using human IFN-γ, IL-2, and granzyme B (GzmB) antibodies (Mabtech). Briefly, MSIPS4W plates (Millipore) were activated with 35% ethanol and coated with primary antibody: 10 μg/mL for IFN-γ (1-D1K), 10 μg/mL for IL-2 (MT2A91/2C95), and 15 μg/mL for GzmB (MT28). Plates were incubated overnight at 4°C, washed with PBS, and blocked with R10 for 1 hour at 37°C. PBMCs were thawed and rested for at least 2 hours prior to plating. Plates were washed and PBMCs were plated at 2 x 10^5^ cells/well. Peptides were added at a final concentration of 1 µg/mL/peptide. Plates were incubated at 37°C for 24 hours (IFN-γ and IL-2) or 48 hours (GzmB). After incubation, plates were washed with PBS containing 0.05% Tween-20 (BioShop) before the addition of human IFN-γ (7-B6-1), IL-2 (MT8G10), and/or GzmB (MT8610) biotinylated secondary antibodies. For dual colour assays, alkaline phosphatase (ALP) conjugated to IFN-γ secondary antibodies were used with other biotinylated secondary antibodies targeting another cytokine. Streptavidin-ALP and/or streptavidin-horseradish peroxidase (HRP) was added after washing and spots were developed using Vector Blue for ALP and/or Vector NovaRED for HRP (Vector Laboratories). All conditions were done in duplicate unless otherwise stated. Spots were quantified with an ImmunoSpot S3 Analyzer (Cellular Technology Limited). For data analysis, mean spots of the negative control wells (DMSO) were subtracted from all peptide-stimulated wells before results were normalized to spot forming cells per million PBMC (SFC/10^6^). Results were considered positive if wells were above the higher of the two thresholds: ≥20 SFC/10^6^ PBMC or twice the mean value of negative control wells.

### Generation of Immortalized B Cell Lines

B cell lines (BCL) were generated using PBMCs and immortalized with Epstein-Barr virus (EBV). Frozen PBMCs were thawed and incubated with 2.5 mL of supernatant from B95-8 cell lines for 2 hours in a 37°C water bath. 1 μg/mL of cyclosporin A in 5 mL of R10 was then added to the cell suspension and incubated in a T25 flask for 3 weeks at 37°C. B cells that were difficult to immortalize in this manner were first purified using a human B cell isolation kit (STEMCELL), then incubated with B95-8 supernatant.

### Generation of SARS-CoV-2-Specific CD4+/CD8+ T Cell Clones and Lines

Frozen PBMCs were thawed and first depleted using CD4+ or CD8+ positive selection kits (STEMCELL). Depleted PBMCs were incubated at 37°C overnight with the peptide of interest at 10 µg/mL in R5 media that contains 5% human serum instead (Wisent). After overnight incubation, SARS-CoV-2-peptide-specific CD4+ T cells were selected using an IL-2 secretion kit (Miltenyi) while SARS-CoV-2-peptide-specific CD8+ T cells were obtained using an IFN-γ secretion kit according to the manufacturer’s instructions (Miltenyi). Enriched T cells were resuspended in R10-Max media that contains 2 mM Glutamax instead (Gibco). T cells were plated in 96-well plates at limiting dilution. T cells were then co-cultured with irradiated feeder cells containing allogeneic PBMC and BCL, 50 U/mL IL-2 (R&D Systems), 10 ng/mL IL-15 (R&D Systems), and 25 ng/mL mouse anti-human CD3 antibody (UCHT1; BD Bioscience). T cells were incubated at 37°C for 4-5 weeks and fed biweekly with R10-Max and 50 U/mL IL-2. Next, IFN-γ ELISpot was used to screen the potential T cell clones for specificity against the peptide of interest (1 μg/mL) in the presence of autologous BCL (1 x 10^3^ cells/well). For T cell lines, PBMCs were pulsed with 50 µg/mL of the target peptide for 1 hour at 37°C, diluted to 1 µg/mL of peptide, and plated in 12-well plates at 2 x 10^6^ cells/mL. After 24 hours incubation at 37°C, 50 U/mL IL-2, and 25 ng/mL IL-7 were added. PBMCs were kept in culture for 3-5 weeks and stimulated weekly with 1 µg/mL of target peptide, 50 U/mL IL-2, and 25 ng/mL IL-7. IFN-γ ELISpot was then used to test the specificity of potential T cell lines against the target peptide (1 μg/mL) in the presence of autologous BCL (1 x 10^3^ cells/well). The T cell compositions of clones and lines that elicited positive IFN-γ ELISpot responses were then examined via flow cytometry.

### HLA Restriction of T Cell Clones and Lines

To determine HLA I-restricted epitopes from CD8+ T cells, a B cell panel containing autologous and allogeneic BCL was used. BCL were pulsed with the specific peptide for 1 hour at 10 µg/mL, washed, then co-cultured with CD8+ T cell clones in an IFN-γ ELISpot. To determine HLA II-restricted epitopes from CD4+ T cell clones or lines, autologous BCL were blocked with 10 µg/mL of either anti-HLA-A/B/C (BD Biosciences), anti-HLA-DP (Abcam), anti-HLA-DQ (BioLegend), anti-HLA-DR (BioLegend), or anti-HLA-DP/DQ/DR (BioLegend) antibodies for 30 minutes, then pulsed with the specific peptide for 1 hour at 10 µg/mL. Pulsed BCL (1 x 10^3^ cells/well) were then washed 3 times with R10, mixed with CD4+ T cells (1-2 x 10^4^ cells/well), and tested in IFN-γ ELISpot assay in the presence of 10 µg/mL of anti-HLA antibodies.

### CFSE T Cell Proliferation Assay

CFSE T cell proliferation assay was performed as previously described (20). Briefly, PBMCs obtained from various time points were pre-labelled with 5 µM of carboxyfluorescein diacetate succinimidyl ester (CFSE; Thermo Fisher Scientific) in PBS with 2.5% FBS for 8 minutes in 37°C water bath. CFSE-labelled cells were then resuspended in R10 media supplemented with recombinant IL-2 (R&D system) and 2-mercaptoethanol (Themo Fisher Scientific) before being plated in 96-well U-bottom plates at 4 x 10^5^ cells/well. PBMCs were then pre-stimulated with 0.1 µg of N, E, M, and S (RBD and TM) master peptide pools, DMSO (Negative control), or SEB (Positive control) and incubated at 37°C for 5 days. On day 6, PBMCs were re-stimulated with 1 µg/mL of master peptide pools in the presence of brefeldin A (GolgiPlug; BD Bioscience) and monensin (GolgiStop; BD Bioscience) for 24 hours. On day 7, PBMCs were first stained with LIVE/DEAD™ fixable blue dead cell stain (Thermo Fisher Scientific) and incubated with Fc receptor blocking solution (Human TruStain FcX^TM^; BioLegend) before surface staining with fluorochrome-conjugated antibodies to CD3 (APC-Cy7: Clone SK7), CD4 (PerCP-Cy5.5: Clone SK3), and CD8 (PE: Clone HIT8a). Cells were then fixed using BD Cytofix/Cytoperm and permeabilized using BD Perm/Wash according to the manufacturer’s protocol before staining with anti-IFN-γ (APC: Clone 4S.B3), anti-GzmB (BV421: Clone GB11), and anti-IL-2 (BV711: Clone MQ1-17H1). All antibodies were purchased from BD Bioscience. Samples were then acquired on the BD Fortessa-X20. Net master peptide pool-induced CFSE_Low_ responses were calculated as the percentage of master peptide pools with reduced CFSE fluorescence minus the percentage of control (DMSO) stimulated cells with reduced CFSE fluorescence.

### Cytotoxic T Lymphocyte (CTL) Killing Assay

CTL killing assays were performed as previously described with minor modifications (21). Autologous BCL were used as target cells and pre-incubated with 40 ng/mL IFN-γ (R&D System) for 18 hours at 37°C. To differentiate peptide-pulsed and non-peptide-pulsed target cell populations, BCL were stained with 0.02 µM CFSE (CFSE_Low_) or 0.2 µM CFSE (CFSE_High_) for 15 minutes at 37°C, respectively. CFSE_Low_ BCL were then pulsed with the target peptide at 5 µg/mL for 45 minutes at 37°C. Both CFSE_Low_ and CFSE_High_-labelled BCL were washed twice with warm R10 prior to mixing at 1:1 ratio and resuspended to 2 x 10^5^ cells/mL for plating. For effector cells, T cell clones were washed three times with warm R10 to remove any excess cytokines and serially diluted from 32:1 to 0.5:1 (Effector:Target Ratio). Both target and effector cells were then plated to 96-well U-bottom plate and incubated for 6 hours at 37°C. Cells were then stained with LIVE/DEAD™ fixable near IR cell stain (Thermo Fisher Scientific) and incubated with Fc receptor blocking solution (Human TruStain FcX^TM^; BioLegend) before surface staining with fluorochrome-conjugated antibodies to CD4 (BV711: Clone SK3) and CD8 (PE: Clone HIT8a). In a separate experiment, intracellular cytokines secreted by the T cells clones were determined as previously described by using anti-IFN-γ (APC: Clone 4S.B3), and anti-GzmB (BV421: Clone GB11. All antibodies were purchased from BD Bioscience. Samples were then acquired on the BD LSR Fortessa.

### Experimental Software and Statistical Analysis

All flow cytometry data were analyzed using FlowJo 10.8.0 software (Treestar, Ashland, OR, USA). Prism 9.0 (GraphPad, San Diego, CA, USA) was used to perform statistical and graphical analyses.

### Within-Host Model for SARS-CoV-2 Infection and Subsequent Immune Response

To model the initial infection and subsequent production of infectious virions, we used a target cell-limited model with an eclipse phase **(Eq. 1a-d, full model depicted and described in Supplemental Figure S3)** similar to that successfully used to model Influenza A (22, 23), as well as SARS-CoV-2 (24). Our full model is given by:

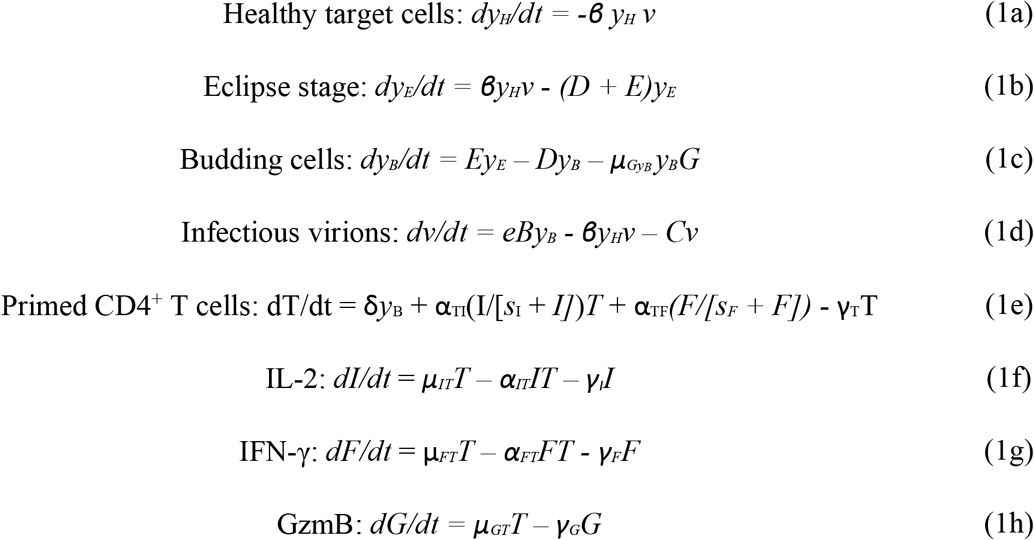

### Within-Host Model Parameter Estimation and Fitting Assessment

All fits to clinical data using our model (Equation 1) were performed in Monolix (Version 2020R1) using non-linear mixed-effects models. Individual parameters **(Tables S2 and S3)** for each data set are determined by the maximum likelihood estimator Stochastic Approximation Expectation– Maximization (SAEM), and all fits met the standard convergence criteria (complete likelihood estimator). As shown in **Table S2**, many parameters from Eq.1 are fit for this study. However, parameters from Eq.1a-d are chosen based on previous studies on SARS-CoV-2.

## Results

### Clinical Data of Participants

Clinical data of SARS-CoV-2 infected subjects can be found in **Table 1**. 23 subjects were sampled at the time of presentation, either during acute infection and/or shortly after convalescence, ranging between 7-160 days post symptom onset (PSO). However, only 21 subjects were available for subsequent blood draws to understand long-term persistence of immune memory to SARS-CoV-2 (2-7 blood draws up to 398 days PSO). All subjects were infected with the ancestral circulating Wuhan strain. Disease severity ranged from mild (non-hospitalized), moderate (hospitalized not ICU) to severe (ICU admission). Out of the 23 subjects studied, 9 were female (39%) and 14 were male (61%), with an average age of 50 ranging from 23-72 years; 14 had mild (61%), 6 had moderate (26%), and 3 had severe (13%) COVID-19 disease. Two subjects (OM8100 and OM8123) received one dose of the Pfizer BNT162b2 COVID-19 mRNA vaccine in between study visits. Another subject (OM8126) received two doses of the Pfizer vaccine in between study visits. Healthy controls subjects included individuals who had blood drawn either pre-COVID (before January 2020) or were asymptomatic with no history of viral illness and had negative SARS-CoV-2 serology. Three individuals who were hospitalized were sampled intensively at 7-10 day intervals during and after their hospital admission for 6 weeks. The remainder of individuals were sampled during their convalescence after most symptoms had resolved.

**Table 1:** Clinical Summary of SARS-CoV-2 Infected Subjects

| **Disease Severity** | Asymptomatic/Mild | Moderate | Severe | Total |
| --- | --- | --- | --- | --- |
| **No. of Participants (%)** | 14 (61%) | 6 (26%) | 3 (13%) | 23 |
| **Male/Female (n)** | 7/7 | 4/2 | 3/0 | 14/9 |
| **Age (Year)**<br>**Mean ± SD** | 47 ± 14 | 59 ± 9 | 49 ± 6 | 50 ± 13 |
| **Sample Collection**<br>**Range (First Visit)** | 9-160 Days PSO | 7-78 Days PSO | 29-34 Days PSO | Median: 36 Days PSO |
| **Sample Collection**<br>**Range (Last Visit)** | 191-392 Days PSO | 169-360 Days PSO | 377-398 Days PSO | Median: 336 Days PSO |

### Cytokine Effector T Cell Responses at Presentation

Effector T cell responses that include IFN-γ (25), IL-2 (26), and GzmB (27) production are important in viral control of a Th1-mediated anti-viral response. Cytokine ELISpot assay is a highly sensitive methodology to detect low frequency viral antigen-specific responses in *ex vivo* PBMC during or after virus infections (28). We measured SARS-CoV-2 antigen-specific IFN-γ, IL-2, and GzmB responses at presentation by cytokine ELISpot assay. We measured responses to the structural proteins of SARS-CoV-2 that included Spike (S1 and S2), Membrane (M), Nucleocapsid (N), and Envelope (E) **(Figure 1a**). In addition, we included a peptide pool only spanning Spike receptor binding domain and transmembrane domains (S-RBD and S-TM) in certain experiments to further evaluate T cell responses to this region. A representative example of cytokine ELISpot assays performed on *ex vivo* PBMC are shown in **Figure 1a.**

**Figure 1:**
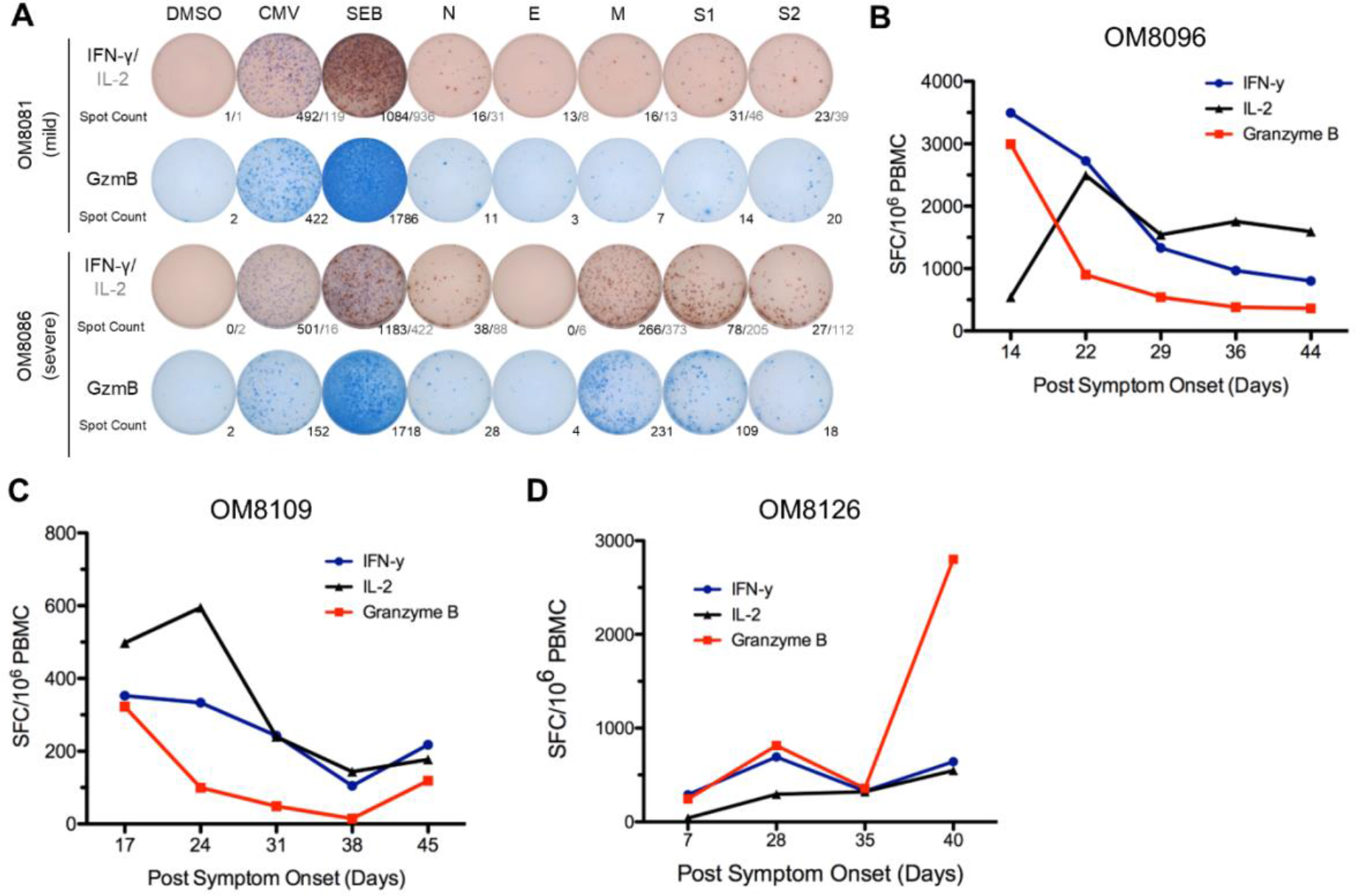
Stronger overall T cell cytokine ELISpot responses detected in severe patients and overall ELISpot responses of acute patients in the first six weeks PSO. **(A)** Representative cytokine ELISpot responses from individuals with mild or severe severe illness against SARS-CoV-2 structural proteins (N, E, M, S1 & S2), with DMSO as negative control and CMV and SEB as positive controls. Numbers indicate the number of spot forming cells for IFN-γ/IL-2 or GzmB. **(B)-(D)** Three subjects with acute moderate COVID-19 infection were followed weekly shortly after symptom onset for up to 6 weeks PSO. Total additive response to the four SARS-CoV-2 structural proteins was measured by ELISpot for each cytokine.

### T Cell Immune Responses During Acute Infection in Hospitalized Individuals with Moderate Illness

We studied 3 individuals during the acute stage of SARS-CoV-2 infection all who were hospitalized (moderate disease), over a 6-week period, from day 7 to 44 days PSO and assessed cytokine ELISpot responses over this time interval at about 7-10 day intervals. Total effector cytokine SARS-CoV-2 specific responses are depicted in **Figure 1 b-d** for each individual. Although all three individuals had moderate disease and were discharged from hospital, we find considerable dynamic fluctuations in their early T cell cytokine responses to SARS-CoV-2 antigens. Cytokine antigen-specific responses were detectable at all time points. We tended to see an early peak of T cell cytokine responses in the first 22-28 days, followed by a leveling of the response afterwards.

### Memory SARS-CoV-2 T Cell Responses in Convalescent Patients

To examine the duration and the strength of the T cell responses against SARS-CoV-2 structural proteins (N, E, M, S1 - 5’ region of spike with RBD, S2 - 3’ region of spike after RBD), 21 convalescent (most symptoms resolved) subjects (13 Mild, 6 Moderate, and 2 Severe) were examined longitudinally via *ex vivo* cytokine ELISpot, where the median time days PSO of first blood draw was 38 days (range 7-160 days PSO). In the majority of subjects, the first time point sampled we usually observed the most potent cytokine responses. A summary of all cytokine responses of all individuals during their maximal response during convalescence is depicted in **Figure 2**. As for the most frequent T cell targets of SARS-CoV-2 structural proteins, 7/21 (33%) subjects responded to N most frequently, followed by S2 with 6/21 (29%) subjects, S1 with 5/21 (24%) subjects, and M with 4/21 (19%) subjects. No responses to E protein were observed. The greatest frequency of cytokine producing cells were found in those with moderate and severe disease compared to mild disease. Total *ex vivo* SARS-CoV-2-specific frequencies for severe/moderate disease subjects versus mild disease were for IFN-γ, 852 versus 204 SFC/10^6^ PBMC (p = 0.02), for IL-2, 1358 versus 191 SFC/10^6^ PBMC (p < 0.001), and for GzmB, 1030 versus 331 SFC/10^6^ PBMC (p = 0.023), respectively; thus, indicating that those with severe/moderate disease had higher frequencies of cytokine-producing SARS-CoV-2-specific T cells than those who suffered mild disease during convalescence. Overall, GzmB cytokine responses from *ex vivo* T cells were greater than IL-2, which were greater than IFN-γ. Among the 21 participants, 11 exhibited GzmB responses as their strongest cytokine response, which were against mainly N and S1/S2. 7 participants possessed IL-2 responses as their strongest response, which were mainly against M and S1/S2. Only 3 subjects exhibited IFN-γ as their strongest responses, which targeted mainly N and S1/S2. The majority of the cytokine response was contributed by CD4+ T cells, since CD4+ T cell depletion of PBMC resulted in reduction of all cytokine responses by 90% (range 70-95%, see **Supplemental Figure S1**, and data not shown). In order to further understand the intensity of SARS-CoV-2-specific T cell responses in convalescent individuals, we also looked at uninfected individuals (pre-2020 and asymptomatic SARS-CoV-2 seronegative individuals) since pre-existing immunity to seasonal alpha and beta coronaviruses may impart cross-reactive immune responses to SARS-CoV-2 antigens, as recently demonstrated (12, 29, 30). We find that at least 50% of uninfected and pre-COVID-19 individuals can demonstrate cross-reactive immune responses to SARS-CoV-2 proteins, **(Supplemental Figure S2)**, with 6/11 SARS-CoV-2 negative individuals having detectable IFN-γ responses, which were generally observed at lower frequencies than convalescent individuals.

**Figure 2:**
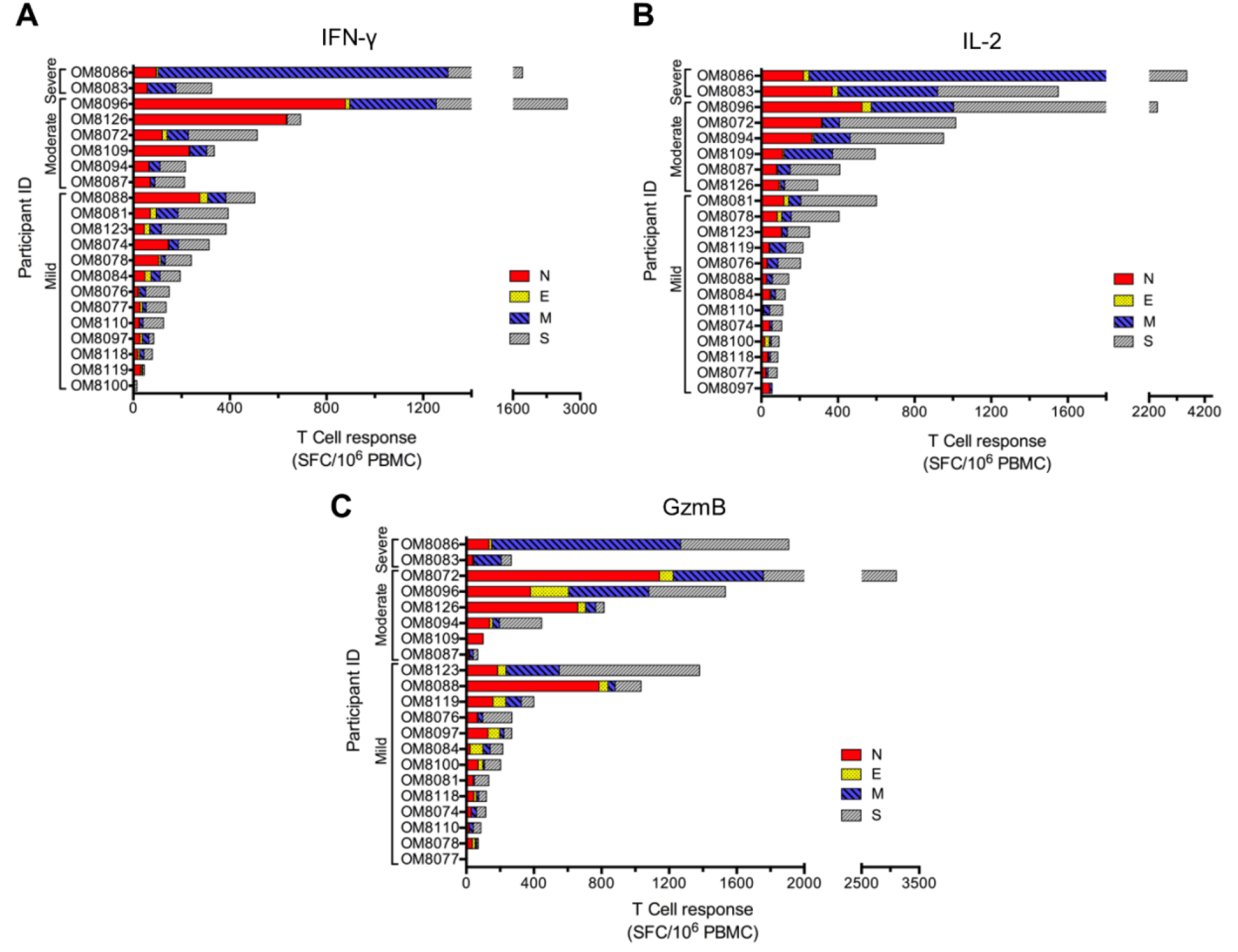
Summary of peak T cell responses of 21 subjects during convalescence. Overall peak ELISpot responses against SARS-CoV-2 structural proteins (N, E, M, S1 & S2) during convalescence of mild (n = 13), moderate (n = 6), and severe (n = 2) subjects. Spot numbers of overall responses were combined from each of the structural protein peptide pools and peak responses from each subject’s time point were selected for **(A)** IFN-γ, **(B)** IL-2, and **(C)** GzmB.

We followed *ex vivo* cytokine responses over a one year period (range 169 to 398 days PSO). In general, we saw a decline in *ex vivo* SARS-CoV-2 responses to all antigens and cytokines. A representative example of one individual is shown in **Figure 3**. To understand the decay of immune memory, the frequency of low-level SARS-CoV-2 *ex vivo* responses that declined to below 50 SFC/10^6^ PBMC to all antigens combined (N, E, M, and S1 + S2) was 33% for IFN-γ, 14% for IL-2, and 29% for GzmB at a median time point of 336 days PSO. Three individuals also received BNT162b2 vaccine during follow-up and showed variable ELISpot responses post vaccination: one subject (OM8100) showed increased IFN-γ/IL-2/GzmB responses to Spike (S1 + S2), but continued decay of N, M after vaccination; the second subject (OM8123) showed continued decay of responses after vaccination; the third subject (OM8126) showed only increased IFN-γ responses only to S1 but no changes against other proteins (data not shown).

**Figure 3:**
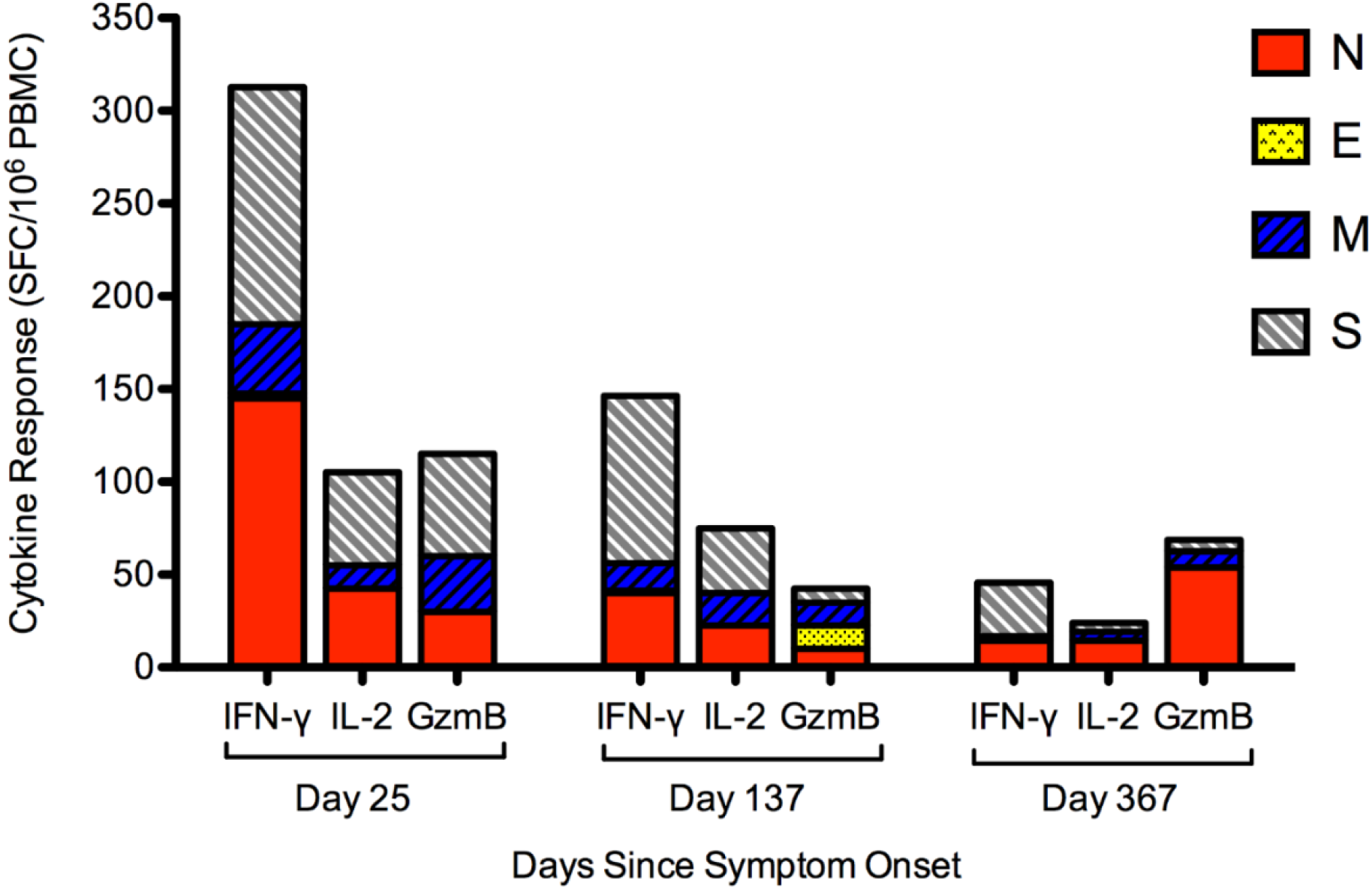
Representative of longitudinal ELISpot data (subject OM8074, mild illness) demonstrating decay of cytokine responses against SARS-CoV-2 structural proteins over time. Overall IFN-γ, IL-2, and GzmB ELISpot responses against SARS-CoV-2 structural proteins (N, E, M, S1 & S2) were measured over a period of one year PSO.

### Modelling of *Ex Vivo* SARS-CoV-2 Specific Immunity Over Time

Using a within-host model (Eq.1), individual fit parameters for the period of data collection for 21 participants are shown for total SARS-CoV-2 IFN-γ/IL-2/GzmB responses in **Figure 4**. The model predicted that the immune response variables peaked at 6 days for IFN-γ, 36 days for IL-2, and 7 days for GzmB. Thus, IFN-γ and GzmB appeared to peak earlier than IL-2. Our model, based on severity of disease, is shown in **Figure 5** detailing the average case-severity predicted responses for disease severity as a function of time. For each immune response variable and for each disease severity, we predicted the average response out to two years PSO. We find that the IFN-γ response to have the highest peak for severe cases with a value of ∼100 SFC/10^6^, followed by moderate (∼90 SFC/10^6^), and then mild (∼31 SFC/10^6^). For IL-2, all case severities peak at the same time, however, severe cases display a peak response twice that of moderate, and moderate disease displays a peak response four-fold higher than compared to mild cases. In contrast, GzmB displays little qualitative severity dependence. We then looked at average cytokine production and degradation rates with disease severity (**Figure 5 d-i**). The model predicts that severe illness is associated with a reduced IFN-γ and GzmB, but increased IL-2 production rates (**Figure 5 d, e, f)**. For instance, mean IL-2 production (μ_ΙΤ_) is found to be 0.0066±0.0010 d^-1^, 0.00685±0.001 d^-1^, and 0.00891±0.00123 d^-1^, for mild, moderate and severe disease, respectively **(Figure 5 e**). Mean IL-2 degradation (γ_1_) is found to be 0.107089±0.015004 d^-1^, 0.0983±0.01608 d^-1^, and 0.0728±0.01303 d^-1^, for mild, moderate and severe, disease respectively. Mean GzmB production was found to be 0.2632±0.117 d^-1^, 0.1623±0.0915 d^-1^, and 0.115±0.0517 d^-1^, for mild, moderate and severe disease, respectively. We also applied our model based on sex **Figure 6.** We find that for IFN-γ, males display faster stimulation and slower degradation rates compared to females, whereas for GzmB, we find that females display faster stimulation and slower degradation than males. For IL-2, stimulation and degradation rates are similar between males and females.

**Figure 4:**
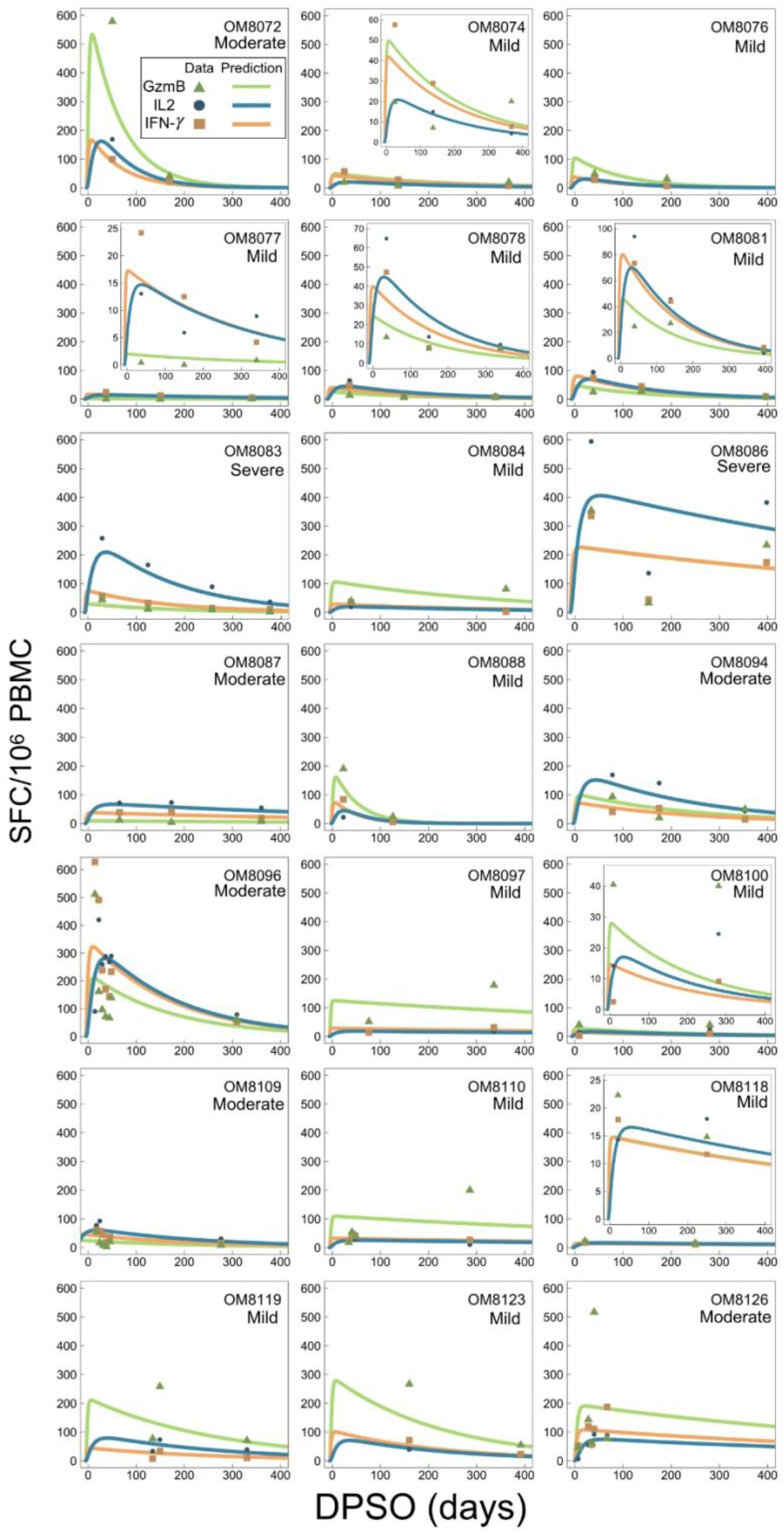
Individual fits as a function of days PSO to the kinetic model (Eq. 1). IL-2, IFN-γ, and GzmB data points are shown as blue circles, orange squares, and green triangles, respectively. Solid lines are fits to model Eq.1, where IL-2, IFN-γ, and GzmB are fit to Eq.1f, Eq.1g, and Eq.1h, respectively.

**Figure 5:**
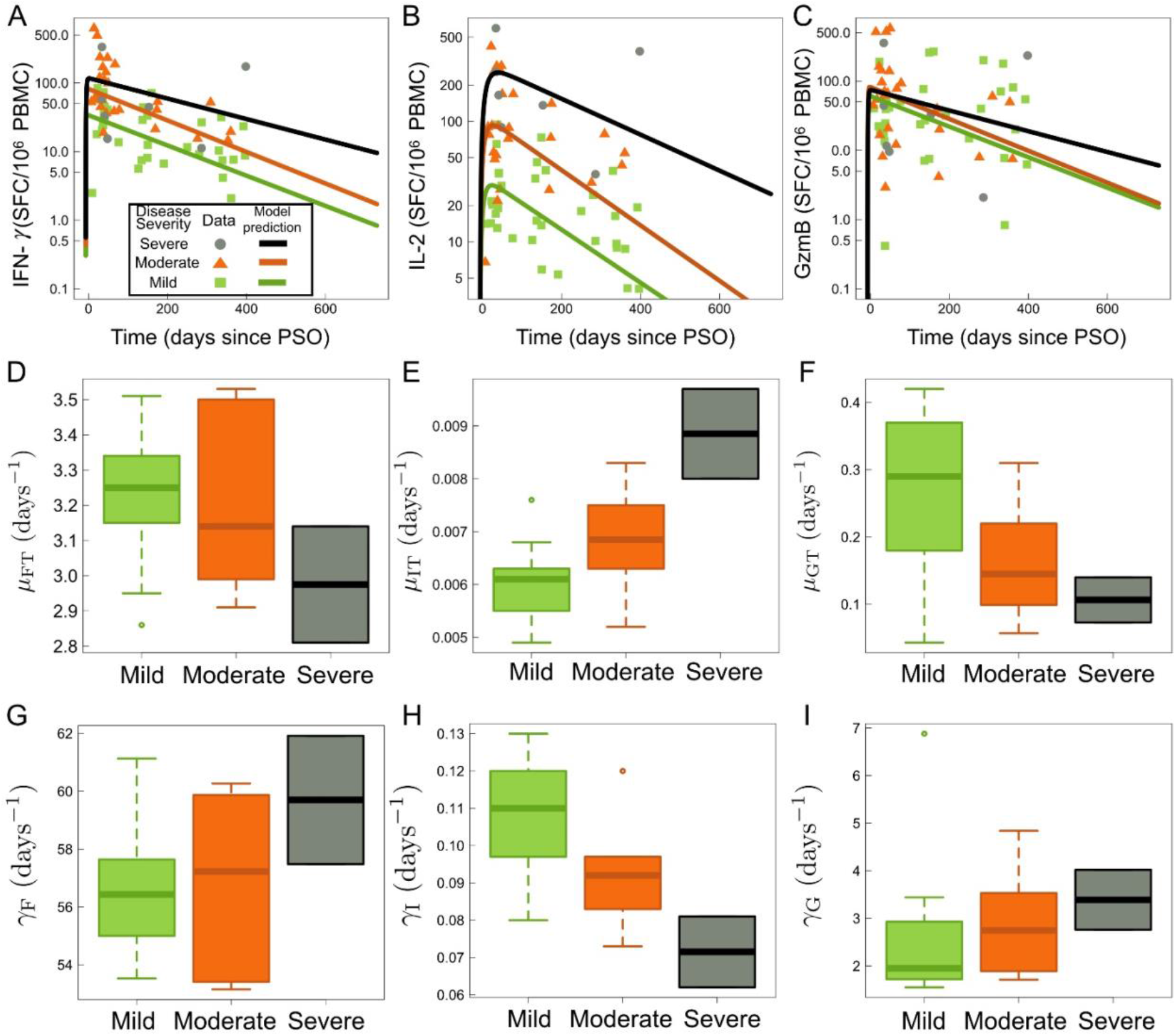
Kinetic model (Eq.1) results sorted by severity. Clinical data in panels **(A)-(C)** is separated by case severity: severe (grey circles), moderate (orange triangles), and mild (green squares). **(A)** IFN-γ data and fits to Eq.1g as a function of days since PSO. **(B)** IL-2 data and fits to Eq.1f as a function of days since PSO. **(C)** GzmB data and fits to Eq.1h as a function of days since PSO. Panels **(D)-(I)** detail boxplots of model fitted parameters for Eq.1fgh sorted by case severity. **(D)** IFN-γ stimulation rate by CD4+ T cells, μFT, for mild, moderate and severe. **(E)** IL-2 stimulation rate by CD4+ T cells, μΙΤ, for mild, moderate and severe. **(F)** GzmB stimulation rate by CD4+ T cells, μGT, for mild, moderate and severe. **(G)** IFN-γ degradation rate, γF, for mild, moderate and severe. **(H)** IL-2 degradation rate, γI, for mild, moderate and severe. **(I)** GzmB degradation rate, γG, for mild, moderate and severe.

**Figure 6:**
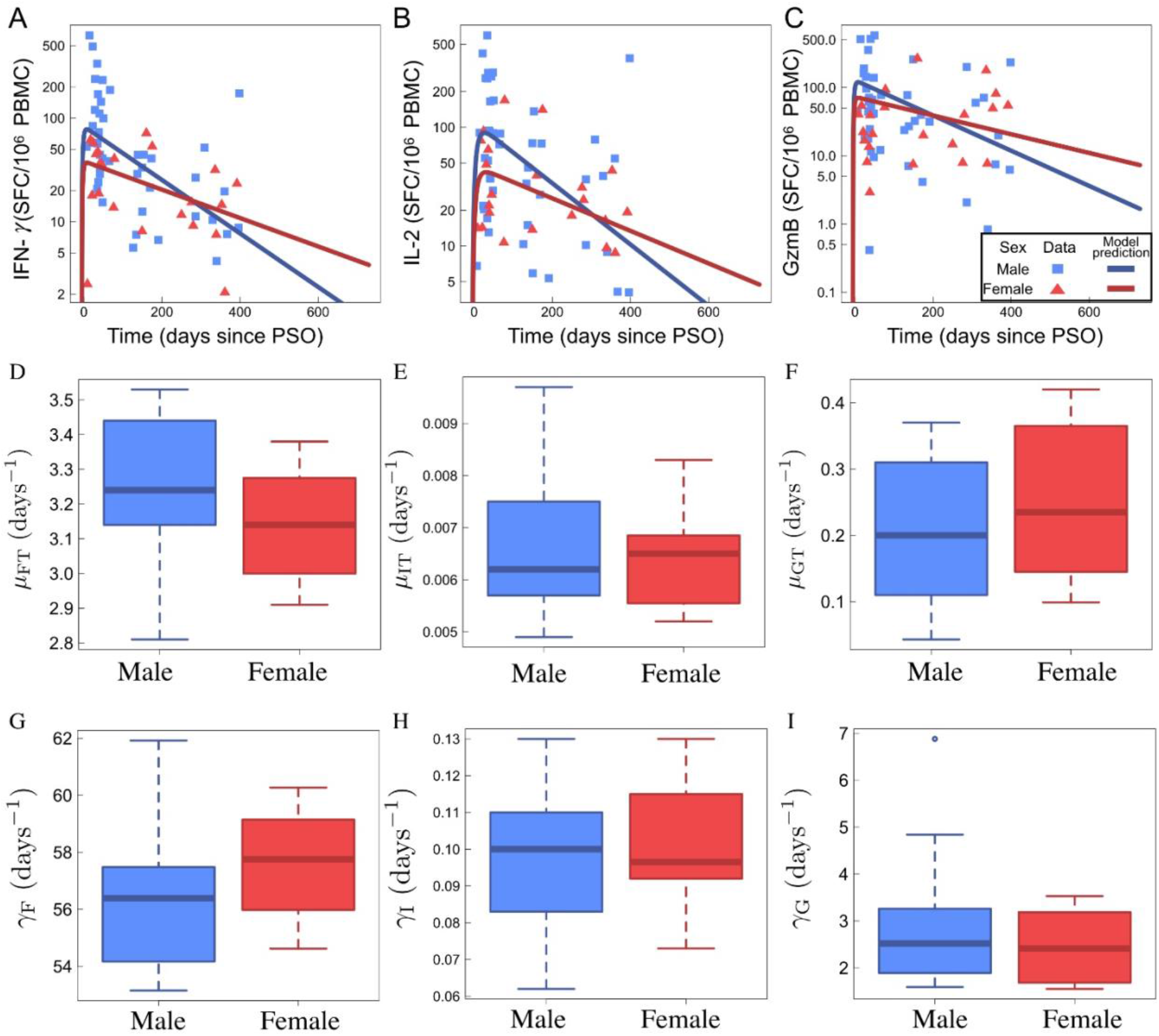
Kinetic model (Eq.1) results sorted by sex. Clinical data in panels **(A)-(C)** is separated by males (blue squares) and females (red triangles), while model fits are shown as blue and red solid lines. **(A)** IFN-γ data and fits to Eq.1g as a function of days since PSO. **(B)** IL-2 data and fits to Eq.1f as a function of days since PSO. **(C)** GzmB data and fits to Eq.1h as a function of days since PSO. Panels **(D)-(I)** detail boxplots of model fitted parameters for Eq.1fgh classified by sex. **(D)** IFN-γ stimulation rate by CD4+ T cells, μFT, for male and female patients. **(E)** IL-2 stimulation rate by CD4+ T cells, μΙΤ, for male and female patients. **(F)** GzmB stimulation rate by CD4+ T cells, μGT, for male and female patients. **(G)** IFN-γ degradation rate, γF, for male and female patients. **(H)** IL-2 degradation rate, γI, for male and female patients. **(I)** GzmB degradation rate, γG, for male and female patients.

Based on the cytokine and GzmB responses, we are also able to estimate the within-host CD4+ T cell half-life from the model Eq.1e. Across all individuals, we find an average CD4+ T cell decay, γT, of 0.005 d^-1^. Where we have assumed single exponential decay kinetics for CD4+ T cells, this value translates to an average half-life of 139 days (or ∼4.6 months). Distributions of CD4+ T cell kinetics for male, female, as well as mild, moderate, and severe individuals can be found in **Supplemental Figure S4.** Lastly, **Supplemental Figure S5** contains predictive checks and 90% confidence intervals for all model-fit results.

### Proliferative T Cell Immune Responses

T cell proliferation in response to viral antigens is important for potent effector and memory responses (31). To further determine virus-specific T cell proliferation capacity over one year PSO, five subjects with either severe (OM8083 and OM8086), moderate (OM8087 and OM8126) or mild (OM8119) symptoms were examined using CFSE-based flow cytometry analysis over multiple time points **(Figure 7)**. Notably, all participants had potent T cell proliferative responses to SARS-CoV-2 antigens during the entire period of observation and maintained proliferative capabilities after one year PSO regardless of disease severity. In this study, cross-reactive proliferative responses were also noted in our healthy donors (OM1 and OM922) after stimulation with SARS-CoV-2 peptide master pools stimulations, albeit they tended to be weaker than the infected individuals. The T cell proliferative responses from infected individuals were driven mainly by CD4+ T cells, apart from OM8126, after one year PSO. This is contrary to our SEB positive control, where most of the T cells proliferative response were either driven by CD8+ only or both CD4+ and CD8+ dominant responses, highlighting the potential downregulation of CD8+ T cell responses during SARS-CoV-2 infection. Additionally, IFN-γ, GzmB, and IL-2 secretion capacity of the CFSE_Low_-responding T cells of subjects OM8083 and OM8086 after one year PSO **(Supplemental Figure S6)** were examined via flow cytometry. Both CFSE_Low_-responding CD4+ T cells and CFSE_Low_-responding CD8+ T cells continually expressed high levels of IFN-γ, IFN-γ/GzmB, IFN-γ/IL-2, or GzmB, further suggesting that convalescent patients have developed effective T cells memory response against SARS-CoV-2.

**Figure 7:**
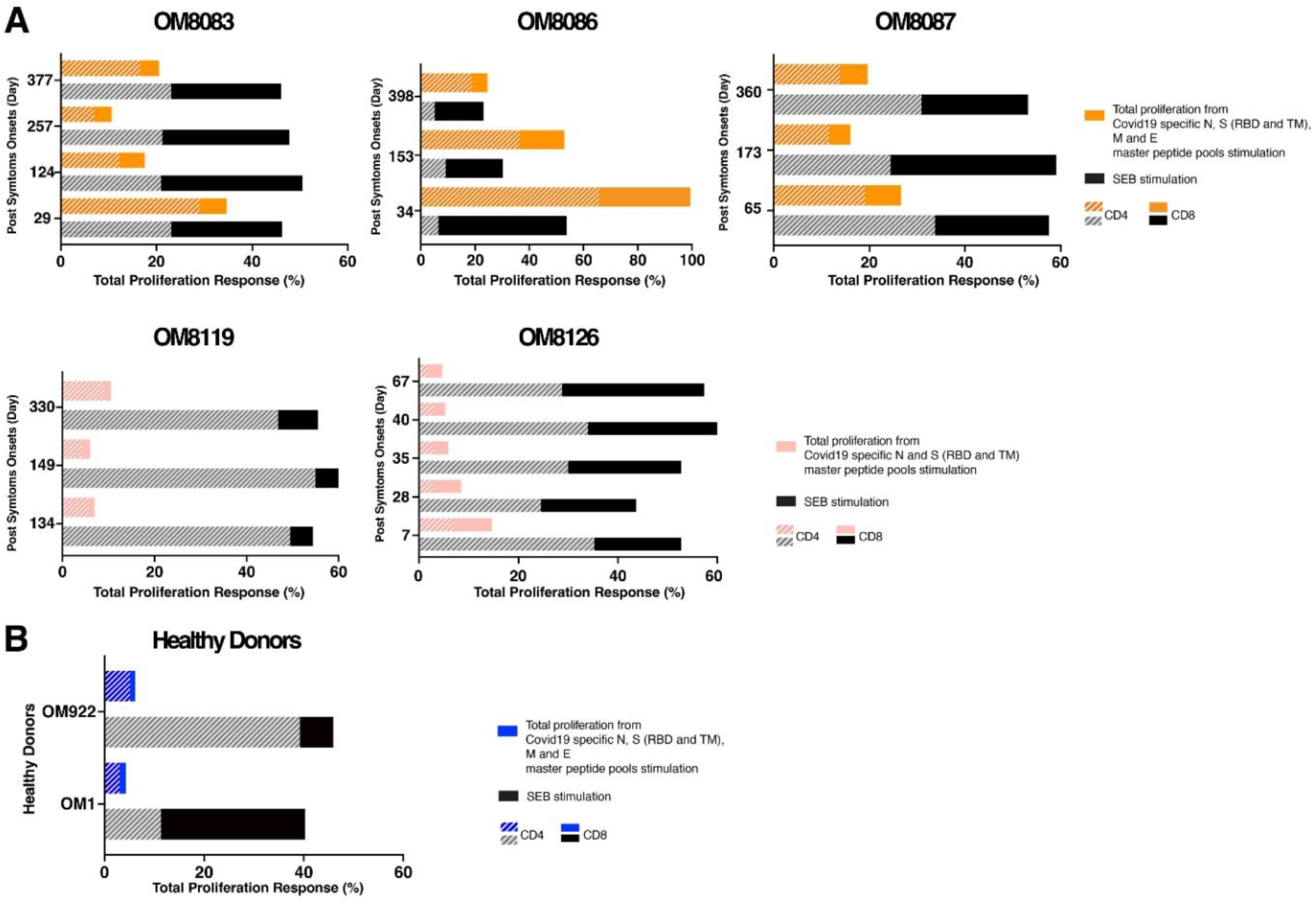
T cell proliferation responses induced by peptide master pools in convalescent COVID-19 subjects over one year PSO. **(A)** Total proliferation from full set peptide master pools stimulations (N, E, M, S-RBD & S-TM) in subjects OM8083, OM8086, and OM8087. **(B)** Total proliferation from full peptide master pools stimulations (N and S-RBD & S-TM) in subjects OM8119 and OM8126. **(C)** Single time point total proliferation response using full set peptide master pools stimulations (N, E, M, S-RBD & S-TM) in two healthy COVID-19 negative donors, OM1 and OM922. Total proliferation was calculated by adding the total of net peptide master pools-induced CFSELow responses from all peptide master pools stimulation. The distribution of CD4+ and CD8+ T cell response in the total proliferation were calculated by averaging CD4+ and CD8+ master pools-induced CFSELow generated at different time points.

### T Cell Epitope Mapping of SARS-CoV-2 Responses

In order to further define antigenic regions targeted by CD4+ and CD8+ T cells, we performed further epitope mapping of IFN-γ responses in 9 individuals (4 Mild, 2 Moderate, and 3 Severe) who demonstrated the strongest *ex vivo* IFN-γ cytokine responses. Representative data are shown in **Figure 8** and summary data are shown in **Table 2**. In some individuals, a substantial frequency of epitope-specific IFN-γ responses could be detected *ex vivo*, varying from 20 SFC/10^6^ upward to ≥4000 SFC/10^6^. As shown in **Table 2**, we were able to define a total of 35 epitopes that belong to ORF1ab (1 epitope), N (13 epitopes), M (14 epitopes), and S (7 epitopes). Shorter amino acid epitopes were not included in the total count since they overlap with the longer amino acid epitopes. Epitopes from M elicited the strongest response in terms of breadth and frequencies in *ex vivo* PBMC, followed by N, S, and ORF1ab. However, we could not fully characterize the S protein epitopes due to unavailability of full S protein peptide matrices at the time of experimentation. Interestingly, no responses to E were observed. Overall, the majority of epitopes that we could define favoured CD4+ T cell responses over CD8+ T cell responses, with greater breadth in individuals with more severe disease **(Table 2)**. Many of the epitopes we identified were previously identified by others (13, 32-51), suggesting common induction of certain epitopes (**see Table 2).** These included one in the ORF1ab region, the alpha and beta coronavirus families cross-reactive N26-N27 region, the N76-N88 region, the M154-156 region, and the M160-163 region **(Table 2)**. Of note is that N26-N27 region was also observed to be cross-reactive in SARS-CoV-1 and the other common cold coronaviruses (HKU1, OC43, NL63, 229E). The remaining regions (N76-88, M154-156, and M160-163) were characterized in SARS-CoV-1 only. Nine additional epitopes that were not studied as extensively have been further characterized in this study **(Table 2)**. We isolated two CD4+ T cell clones from one individual (OM8086). One clone specific for M155 (LRGHLRIAGHHLGRC) was mainly restricted to HLA-DR, although some inhibition (∼40%) was observed in the presence of anti-HLA-DQ antibodies **(Figure 9),** suggesting some promiscuity of this epitope. The other SARS-COV-2 CD4+ T cell clone specific for M156 (LRIAGHHLGRCDIKD) was restricted to HLA-DR only (**Figure 9**). Our findings for M155 and M156 were consistent with previous studies (12, 13, 36, 43, 48, 52). Specifically, two studies have demonstrated that M155 and M156 were mainly restricted by HLA-DRB1*11, which corresponded to the HLA of our subjects (36, 43) (**Supplemental Table S1**). Among the various ORF1ab peptides tested, a response to TTDPSFLGRY was easily detected in 33% of individuals and was determined to be HLA-A*01:01-restricted after testing with a panel of B cell lines (**Figure 9 c)**. This immunodominant epitope and HLA-restriction was previously reported in other studies (32-35). We isolated a CD8+ T cell clone from one individual recognizing an epitope in the M region, RNRFLYIIK (M128-6), that was restricted to HLA-A*30:01 (**Figure 9**). Only *in silico* analyses were conducted for this particular sequence and HLA restriction (47). However, no responses against RNRFLYIIK were observed in two other HLA-A*30:01 patients in our cohort (data not shown). Interestingly, another study had characterized a similar epitope shifted by one amino acid, NRFLYIIKL, to be restricted to HLA-C*07 (34). However, no responses against NRFLYIIKL were detected in three of our HLA-C*07+ subjects (data not shown). Although 4/9 of our subjects that were screened for epitope mapping were HLA-A*02:01 restricted, we could not detect any CD8+ T cell responses that were restricted to this common allele.

**Figure 8:**
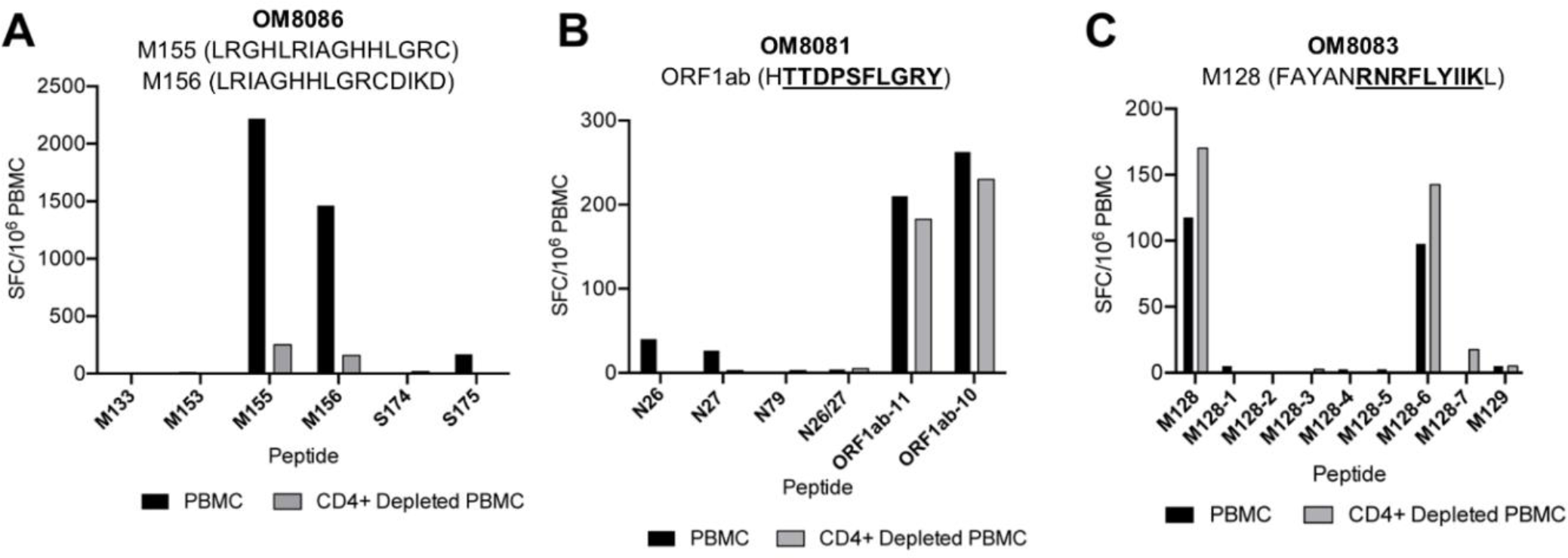
Epitope mapping of SARS-CoV-2 specific T cell responses. Representative graphs of epitope mapping using IFN-γ ELISpot with PBMCs or CD4-depleted PBMCs from three individuals. The strongest peptide per subject is described in the graph header with the minimal epitope underlined where appropriate. **(A)** Individual recovered from severe disease; highly responsive to the 15-mers M155 (LRGHLRIAGHHLGRC) and M156 (LRIAGHHLGRCDIKD) indicating a CD4 epitope. **(B)** Individual recovered from mild disease; highly responsive to the CD8 T cell epitope 10-mer ORF1ab-10 (TTDPSFLGRY). **(C)** Individual recovered from severe disease; highly responsive to the CD8^+^ T cell epitope 9-mer M128-6 (RNRFLYIIK). For the 15-mer M128, 9-mers M128-1 to M128-7 were used to find the minimal epitope.

**Figure 9:**
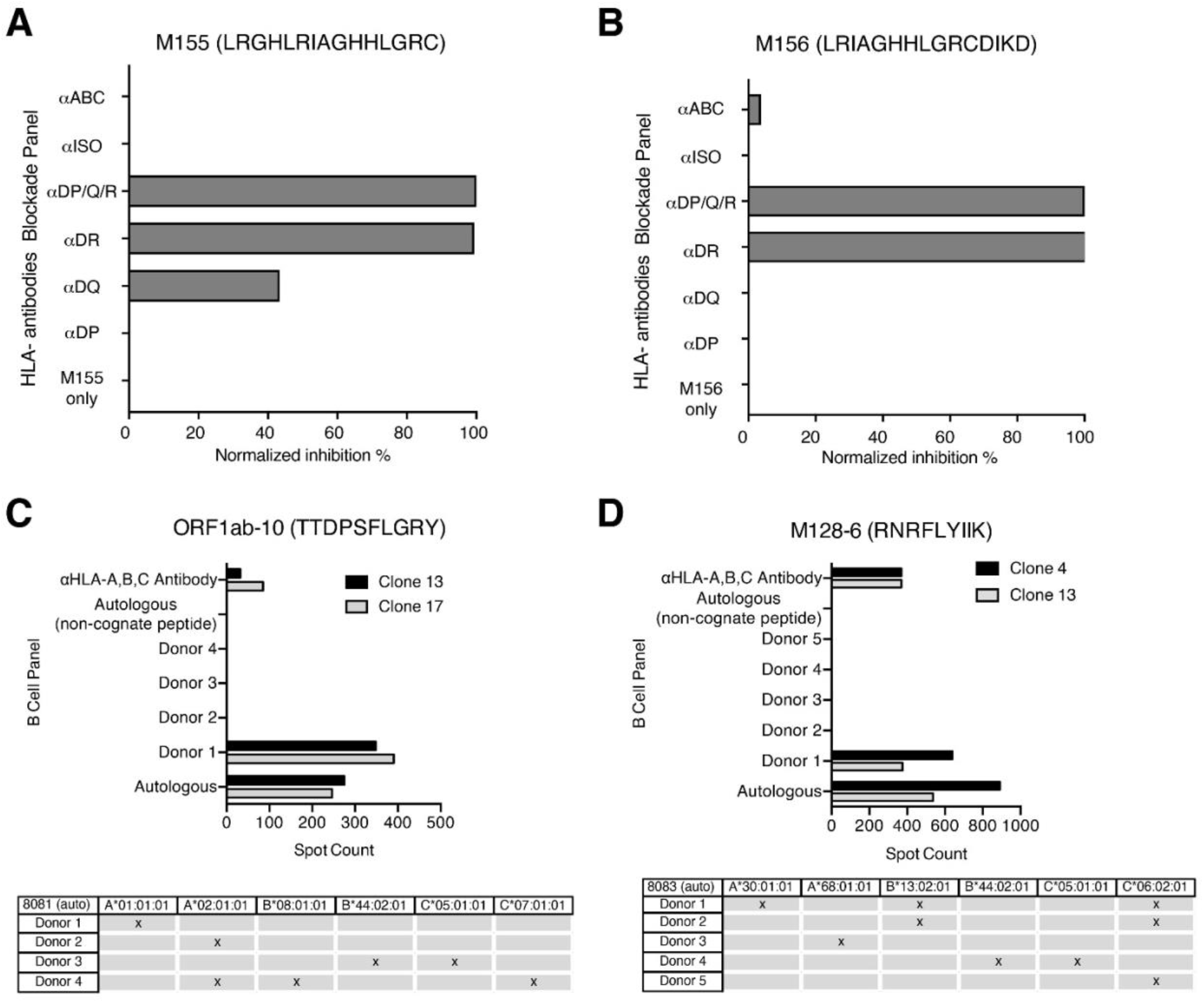
HLA restriction of two CD4^+^ and CD8^+^ epitopes. **(A)-(B)** CD4^+^ T cell epitopes M155 and M156 were HLA-restricted using anti-HLA-II antibodies on CD4^+^ T cell clones stimulated with autologous B cell lines presenting cognate peptide. **(C)-(D)** CD8^+^ epitopes ORF1ab-10 and M128-6 were HLA-restricted using a panel of subject-derived B cell lines. CD8^+^ T cell clones were co-cultured with autologous or allogeneic B cells from the panel pulsed with cognate peptide. All responses here were measured with IFN-γ ELISpot.

**Table 2:**
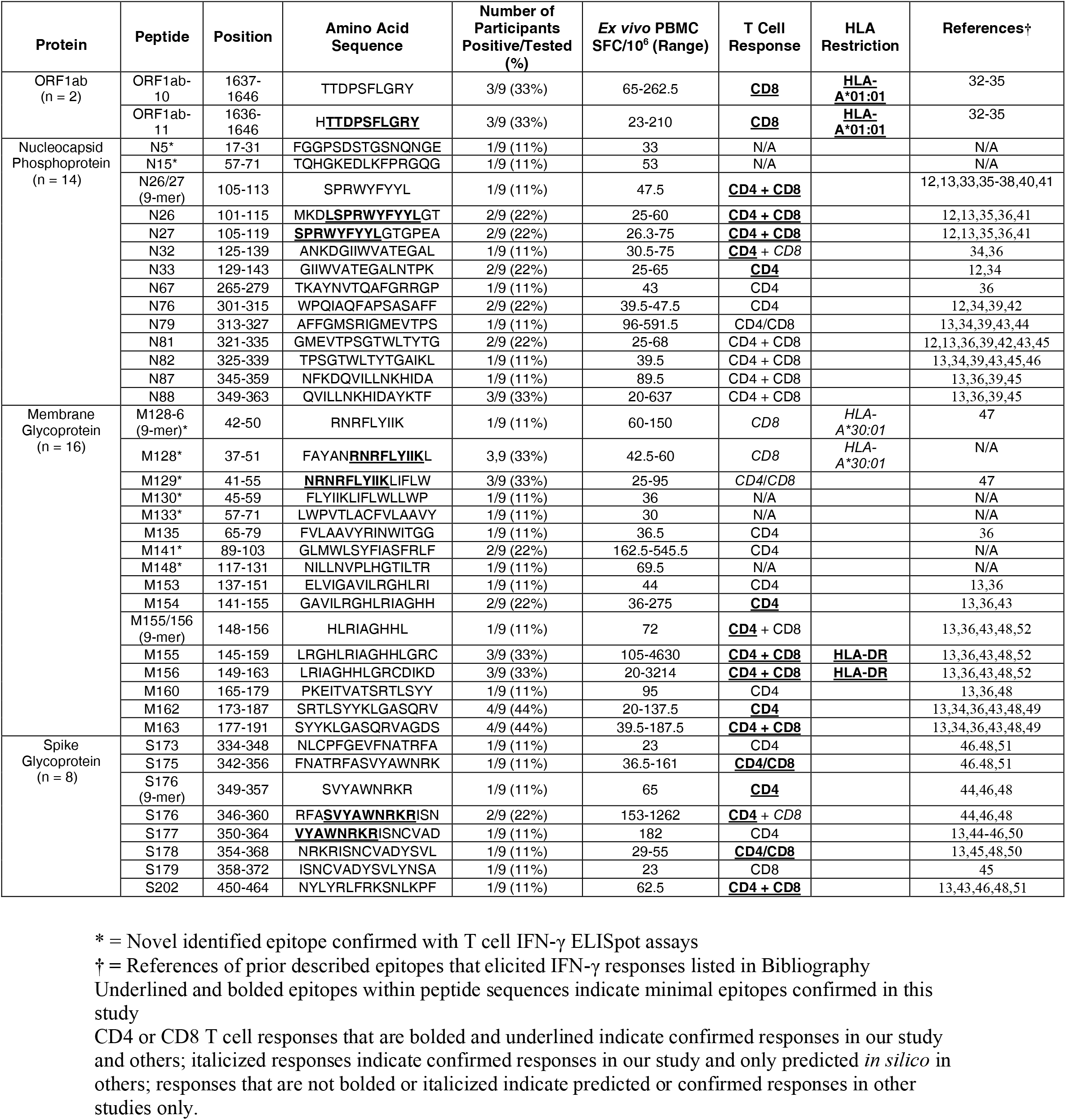
List of Identified T Cell Epitopes

### CD4+ and CD8+ SARS-CoV-2-Specific T Cells Can Recognize and Kill Target Cells

Both SARS-CoV-2-specific CD4+ and CD8+ T cell clones were able to kill peptide-pulsed autologous B cell line target cells (**Figure 10 a, b, c**). Interestingly, SARS-CoV-2 CD8+ T cell clones tended to have enhanced cytotoxic capabilities compared to the SARS-CoV-2 CD4+ T cell clones since they could kill at lower effector:target cell ratios (**Figure 10 b, c**). Both CD4+ and CD8+ T cell clones elicited strong IFN-γ and GzmB responses when stimulated with their specific peptide of interest **(Figure 10 d, e).**

**Figure 10:**
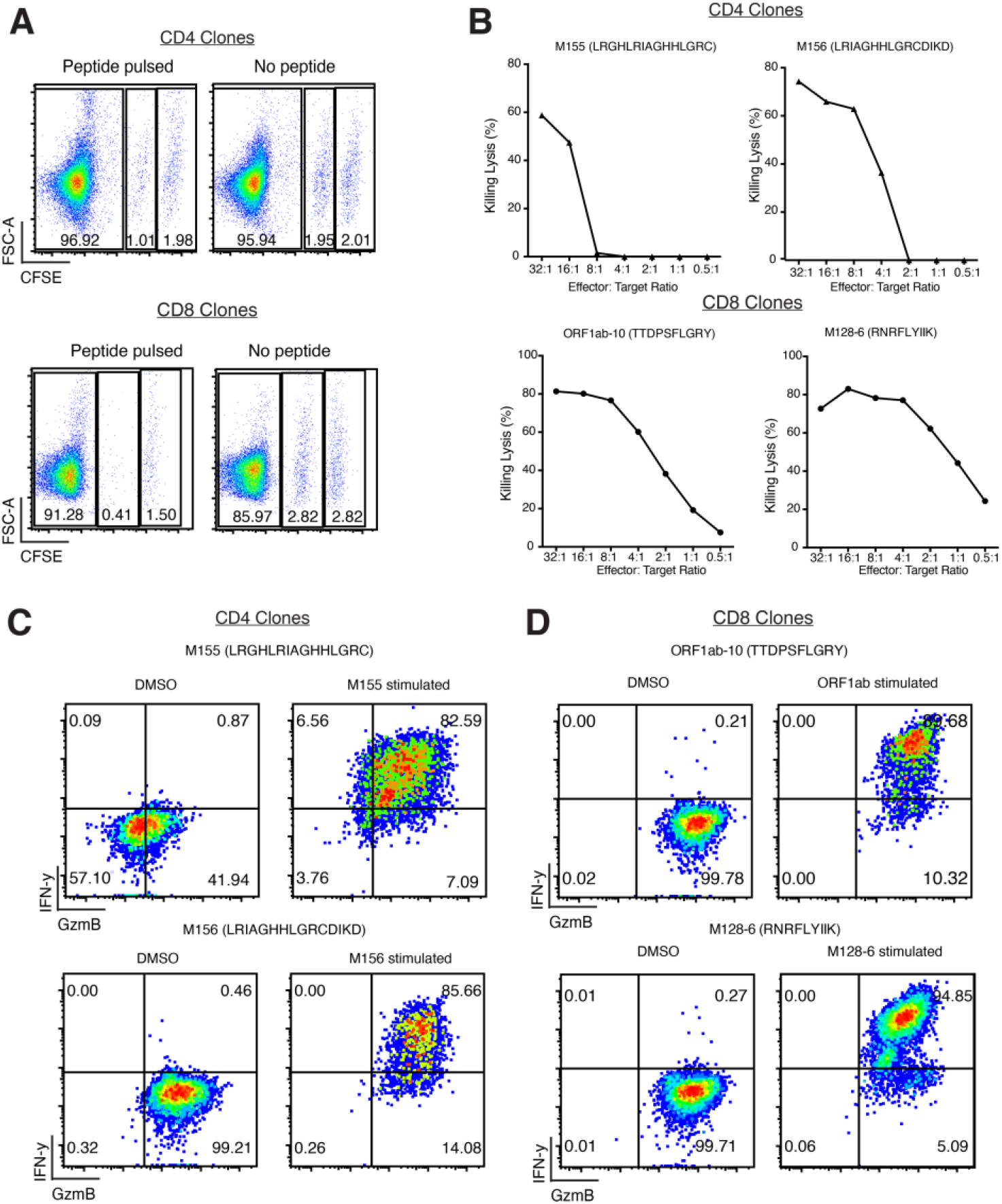
Assessment of recognition and killing capabilities of SARS-CoV-2 Membrane/ORF1ab-specific T cell clones. **(A)** Representative recognition and killing capabilities of SARS-COV-2-specific CD4+ and CD8+ T cell clones against peptide-pulsed, CFSELow target cells. **(B)** Percentage of killing lysis at different effector:target ratios of CD4+ and CD8^+^ T cell clones. **(C)-(D)** Flow cytometry analysis of intracellular cytokines secretion of IFN-γ and GzmB by CD4+ and CD8+ T cell clones in the presence of peptide-pulsed target cells.

## Discussion

The establishment of cellular-mediated immunity is critical against long lasting immunity against most viral infections including coronaviruses. After SARS-CoV-1 infection, antibody-mediated immunity was seen to wane over time (53, 54), however, SARS-CoV-1-specific memory T cells were observed to persist for 6 to 11 years post-infection in several studies, which were at higher frequencies in those with more severe illness (39, 42, 53, 54). In this study, we characterized cytokine producing T cell responses to SARS-CoV-2 structural proteins during acute infection and found potent Th1-focused pluri-cytokine producing responses early and during convalescence in SARS-CoV-2 survivors, with frequencies ranging from 100 to over a 1000 cytokine producing cells per 10^6^ PBMC. These frequencies are similar to those reported by Le Bert et. al. (12) who reported IFN-γ ELISpot against N in SARS-CoV-2 infected individuals. Our data expands these findings to include IL-2, GzmB, and IFN-γ toward all structural proteins. It should be noted that previous estimates describing SARS-CoV-2-specific T cells measured higher frequencies of antigen-specific cells when using *ex vivo* flow cytometric techniques based on activation marker upregulation when exposed to peptide pools (14, 29, 55). However, these frequencies may not necessarily reflect the circulating numbers of antigen-specific cells that are capable of producing cytokines. A number of conclusions can be made from our observations which may in part, be unique to SARS-CoV-2: The SARS-CoV-2 T cell response preferentially induces GzmB and IL-2 over IFN-γ; CD4+ T cell responses make up the majority of this response; T cell responses favoured N, S, and M, but not the E structural proteins; and more severe disease elicited higher memory responses in survivors. The role of GzmB in inducing a potent response is likely important for viral clearance; as previous studies have shown a role of granzymes in control of viral infections in humans, such as acute HIV (27, 56). Although virus infections generally induce very potent CD8+ T cell expansions, this is not observed in our and others’ cohorts for SARS-CoV-2 infection, though it was observed for SARS-CoV-1 (39, 54, 55, 57). The reasons for this are currently, unclear, although a recent study suggests that SARS-CoV-2 ORF8 protein may interfere with MHC class I presentation, thus possibly preventing conventional CD8+ T cell priming (58).

Our data showed a number of features of the kinetics of SARS-CoV-2 T cell immunity. The mathematical modelling results showed early induction of the effector cytokines IFN-γ and GzmB, followed by IL-2 later which is consistent with the effector role of IFN-γ/GzmB in viral clearance and IL-2 to induce long-term memory. Modelling also showed that disease severity induces greater IFN-γ/IL-2, consistent with increased antigen load during moderate and severe disease. The similar production and decay rates of GzmB in mild disease suggests that GzmB may have greater importance at viral clearance, resulting in milder disease course. Our decay rates of T cell cytokine responses were similar to those of Dan et. al. (14), where they observed a half life decay of 3-5 months and we predicted 4.6 months. Our modeling studies also indicated that males and females may differ in IFN-γ and GzmB production in median and variance during infection. Whether these findings help explain the increased mortality rates in men over women to COVID-19 illness will require further study (59, 60). We followed our cohort for at least one year, and encouragingly still find persistence of T cell immunity in the majority of individuals, in particular, SARS-CoV-2-specific IL-2 in 86% and IFN-γ/GzmB in about 70%. Importantly, we were able to find potent proliferative T cell responses that produce cytokines even at the last time points in all individuals studied. These findings indicate that although decay of T cell immunity is observed over one year, one can still induce potent proliferative and effector cytokine responses in T cell obtained one year after infection in most, if not all individuals after restimulation. The role that these responses play in protection against infection and/or disease will require further study and follow up.

SARS-CoV-2 T cell epitope discovery will inform future T cell-based coronavirus vaccines, particularly in the setting of antibody escape variants. Several putative epitopes found in SARS-CoV-1 were, interestingly, observed in SARS-CoV-2. Specifically, the identification of similar epitopes in the N26-N27, N76-N88, and M154-M163 regions in both SARS-CoV-1 and SARS-CoV-2, and other coronavirus families suggest that the common induction of certain epitopes across coronavirus families could be used for vaccine development. We have also described a number of new epitopes that could be incorporated in T cell vaccine designs. For example, the M128-6 epitope (RNRFLYIIK) was a CD8+ HLA*30:01 restricted epitope. In addition, we observed cross-reactive T cell responses in a large number of uninfected individuals which likely reflect previous infection with seasonal coronaviruses. Recent work suggests that these cross-reactive responses may be important in alleviating, rather than exacerbating disease caused by SARS-CoV-2 (61). The identification of additional, novel immunodominant epitopes, such as M128-6 and M141, and the demonstration of recognition and cytotoxic capabilities of CD8+ and CD4+ T cell clones in the M region will be critical for next generation vaccines against potential Spike antibody escape variants. Of the epitopes induced on our subjects, a few mutations are observed in the omicron variant (A63T for peptide M133, and del of 31 for peptide N5 in BA.1, and S371F for peptide S170 in BA.2, see **Table 2**). Further work will determine if these mutations confer escape from the T cell responses. Despite evading CD8+ T cell response induction, we find that CD4+ T cell clones can kill peptide-pulsed target cells, albeit at seemingly lower effector:target cell ratios than the CD8+ T cell clones that we isolated. These features may reflect one reason why SARS-CoV-2 can induce severe disease in a subset of individuals. A combination of a poor CD8^+^ T cell response, with greater killing capacity and an over-exuberant CD4^+^ T cell responses may contribute to disease severity. Thus, further studies will be needed to determine the relative functional avidity of CD4+ vs CD8+ T cells targeting SARS-CoV-2.

In conclusion, we show that SARS-CoV-2-specific memory T cells remain detectable after one year PSO and are capable of proliferating and generating IFN-γ, IL-2, and GzmB responses against SARS-CoV-2 structural proteins, which will likely influence disease course during re-infection of variants and may help explain superior responses to vaccines.

## Author Contributions

JL, RL, CZ, WHK, PS performed experiments;

AE, AJM, JAL, SM, MO recruited subjects

ACG performed SARS-CoV-2 serologies

FYY, MB, PLC, KP processed patient samples

JMH, CSK, MSG, JHKO performed mathematical modeling

JL, MO, CSK, JMH designed the experiments and/or wrote the paper.

## Acknowledgements

The research was funded by grant VR1-172711 from the Canadian Institutes of Health research and the Juan and Stefania fund for COVID-19 and other virus infections, National Research Council of Canada (Pandemic Response Challenge) (JMH, CSK, MSG, HKO), and the Natural Sciences and Engineering Research Council of Canada (JMH).

M.O. receives funding from the Ontario HIV Treatment Network (OHTN), and the Li Ka Shing Knowledge Institute. CSK is funded by an NSERC postdoctoral fellowship. We thank Bhavisha Rathod and the team of the Network Biology Collaborative Centre, Sinai Health, for serological assessment of the samples.

